# Improved methods for protein and single-molecule RNA detection in *C. elegans* embryos

**DOI:** 10.1101/2021.05.07.443170

**Authors:** Dylan M. Parker, Lindsay P. Winkenbach, Annemarie Parker, Sam Boyson, Erin Osborne Nishimura

## Abstract

Visualization of gene products in *Caenorhabditis elegans* has provided insights into the molecular and biological functions of many novel genes in their native contexts. Single-molecule Fluorescence *In Situ* Hybridization (smFISH) and Immunofluorescence (IF) visualize the abundance and localization of mRNAs and proteins, respectively, allowing researchers to elucidate the localization, dynamics, and functions of many genes. Here, we describe several improvements and optimizations to existing IF and smFISH approaches specifically for use in *C. elegans* embryos. We present 1) optimized fixation and permeabilization steps to preserve cellular morphology while maintaining probe and antibody accessibility, 2) a streamlined, in-tube approach that negates freeze-cracking, 3) the smiFISH (single molecule inexpensive FISH) adaptation that reduces cost, 4) an assessment of optimal anti-fade products, and 5) straightforward quantification and data analysis methods. Most importantly, published IF and smFISH protocols have predominantly been mutually exclusive, preventing exploration of relationships between an mRNA and a relevant protein in the same sample. Here, we present methods to combine IF and smFISH protocols in *C. elegans* embryos including an efficient method harnessing nanobodies. Finally, we discuss tricks and tips to help the reader optimize and troubleshoot individual steps in each protocol.

## 1. INTRODUCTION

### 1.1 Microscopic methods for RNA and protein visualization in *C. elegans*

The spatial and temporal patterns of gene expression in *C. elegans* can provide fundamental insights into their function and importance. By querying the abundance and spatial patterning of mRNA and their protein products in whole animals it is possible to gain insight to their transcription and translation, mRNA stability, modification states of protein, developmental regulation, and their functional roles ^1–5^. Visualizing RNA and protein in the same intact animal requires methods that are sensitive, non-perturbative, and, most importantly, compatible with one another. Traditional approaches to visualizing mRNA and protein simultaneously have either relied on the visibility of a GFP-tagged protein to persist under RNA labeling conditions; or they involve combining IF with low resolution FISH protocols. Here, we introduce methods that improve upon existing *in situ* RNA and protein visualization protocols allowing for concurrent imaging of a wide array of proteins and mRNA with state-of-the-art resolution.

The current gold standard for *in situ* single-molecule RNA detection is single-molecule Fluorescence *in situ* hybridization (smFISH). In smFISH, single-molecule RNA visualization occurs by annealing a series of ∼24-48 fluorescently-labeled short antisense oligonucleotide probes to a transcript of interest in fixed animals^6–8^. Annealing multiple fluorescent probes to an RNA produces a discrete, punctate signal for each individual molecule of RNA *in situ*. Labeling each RNA in this manner permits quantification of both the abundance and localization of individual molecules of RNA. Conventional smFISH protocols have successfully characterized RNA expression in *C. elegans*; however, they are challenged by low signal due to poor photostability for some fluorophores and high background^9^ The probes are also costly. We remedy these issues by optimizing the standard smFISH protocol for *C. elegans*, including comparisons of commercial and homemade reagents, rigorous testing of various antifade compounds, and implementation of a recently developed protocol, single molecule inexpensive Fluorscence *In Situ* Hybridization (smiFISH) to reduce cost^10^.

Visualization of endogenous protein expression by immunofluorescence (IF) has also proved to be an indispensible biological tool in *C. elegans*. IF has several benefits in contrast to other protein detection assays. For instance, western blots provide protein abundance and biochemical information but lack any spatial resolution. However, worm embryos pose a challenge for IF experiments due to their strong eggshell and robust permeability barrier^11,12^. Ultimately, this has resulted in adapted protocols requiring harsh fixatives (aldehydes, picric acid), reducing reagents (B-mercaptoethanol, DTT), enzymatic treatments (collagenase), and demanding a high degree of finesse for freeze-crack permeabilization^11,13^. To overcome these challenges, we have adapted strategies for use in the *C. elegans* embryo with comparatively mild chemical treatments allowing antibody penetrance while leaving protein epitopes intact using a simple one-tube protocol.

Perhaps most importantly, we provide a protocol that combines both IF and smFISH in *C. elegans* embryos. While it is sometimes possible to visualize RNA and protein simultaneously with a standard smFISH protocol through the use of fluorescently tagged proteins, tags like GFP can often bleach during fixation. Moreover, conventional methods of smFISH and IF in worms have been challenging to perform in the same sample, resulting in few published protocols. By optimizing the combined protocol, we have co-imaged single-molecules of RNA in conjunction with the proteins they produce *in situ* in whole animals. Our approach is to first perform immunofluorescence followed by smFISH, with key modifications. RNA quality and FISH probe permeability are maintained by using mild fixation conditions and chemical treatments compatible with immunofluorescence while employing RNAse free reagents throughout the protocol. Notably, for some antibody variants, such as nanobodies, a simplified protocol can sometime be utilized. We present the technical details for each protocol individually, in combination, user-friendly ways to analyze the data, standard controls, and some options for troubleshooting. We present several related protocols for the reader to choose between (Figure 1). This includes a comprehensive protocol to perform sample prep, immunofluorescence, smFISH, and slide preparation in series (Figure 1, Protocol 1). Additional protocols also describe smFISH or smiFISH alone (Protocol 2), immunofluorescence alone (Protocol 3), or an alternative simultaneous immunofluorescence/smFISH approach using nanobodies (Protocol 4).

**Figure 1.**
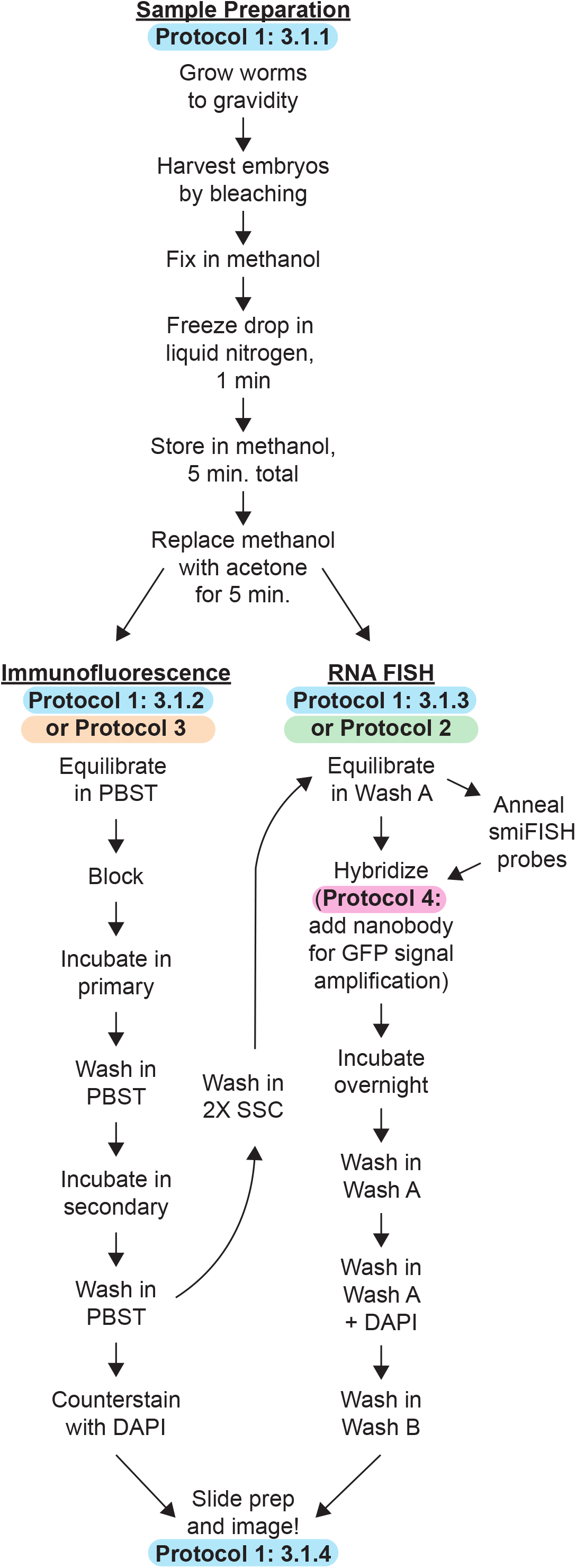
Schematic illustration of IF, FISH, and IF/FISH protocols. An overview illustrating the workflow of the sequential IF/FISH (Protocol 1), RNA FISH (Protocol 2), IF (Protocol 3), and simultaneous IF/FISH (Protocol 4) protocols from sample preparation to slide preparation.

## 2. EXPERIMENTAL DESIGN, CONSIDERATIONS, AND DATA ANALYSIS

### 2.1 Sample Preparation and Fixation

IF and smFISH have been performed using various fixation conditions in *C. elegans* and other model systems. Common fixatives include formaldehyde/formalin or organic solvents such as methanol, ethanol, and acetone. Formaldehyde/formalin acts by creating crosslinked, covalent chemical bonds in the sample, primarily at lysine residues. Formalin can also cause C-T and G-A mutations on DNA sequences as characterized by PCR^14^. Moreover, formaldehyde/formalin-fixation affects tertiary amines in RNA sequences resulting in modification of up to nearly 40 % of As and Cs in formalin-fixed tissues^15^. Due to the high degree of alteration that occurs on nucleic acids, formaldehyde/formalin-fixation is not an ideal fixative for nucleic acid visualization. As an alternative to crosslinking-fixatives, alcohols and other organic solvents have been identified as superior nucleic acid-fixatives^16^. Alcohols and organic solvents, such as ethanol, methanol, and acetone, function by dehydrating clathrate water molecules around proteins and nucleic acids, thus precipitating biological molecules into a fixed state without significant chemical alteration. As with crosslinking fixatives, alcohols and organic solvents have their detriments. These fixatives can disrupt cell membrane structures, cytoplasmic organelles, and soluble cell structural elements such as microtubules^17,18^. However, due to their preservation of nucleic acid composition, they are ideal fixatives for single-molecule RNA detection assays. Further, we have found that short fixations using these types of fixatives allow efficient antibody penetration and do not appear to cause disruption to the protein epitopes we have targeted through IF as some previous studies have shown^19^.

### 2.2 Immunofluorescence

IF has been a staple of *C. elegans* experimentation for decades. As a result, a variety of methods for performing IF have been developed, providing information and protocols for antigen production, peptide coupling, antibody purification, fixation conditions, and protocols related to IF in *C. elegans*^11,20,21^. However, the majority of these methods have focused on the use of larval stages of development, and are not optimized for embyos. Most protocols use some combination of reducing reagents, enzymatic treatments, formaldehyde fixation, and “Freeze-Cracking” mechanical disruption – compressing samples between slides, not to be confused with freeze-cracking of the eggshell in liquid nitrogen – ^13^. Here we present a single-tube protocol requiring no reducing reagents or enzymatic treatments and utilizing a light methanol/acetone fixation and liquid nitrogen cracking to permeabilize the eggshell. We demonstrate this protocol using the anti-PGL-1 antibody K76^20^ (DHSB, Antibody registry ID AB_531836) and the anti-ELT-2 antibody 455-2A4^22^ (DHSB, Antibody Registry ID: AB_2618114) **(FIGURE 2)**.

**Figure 2.**
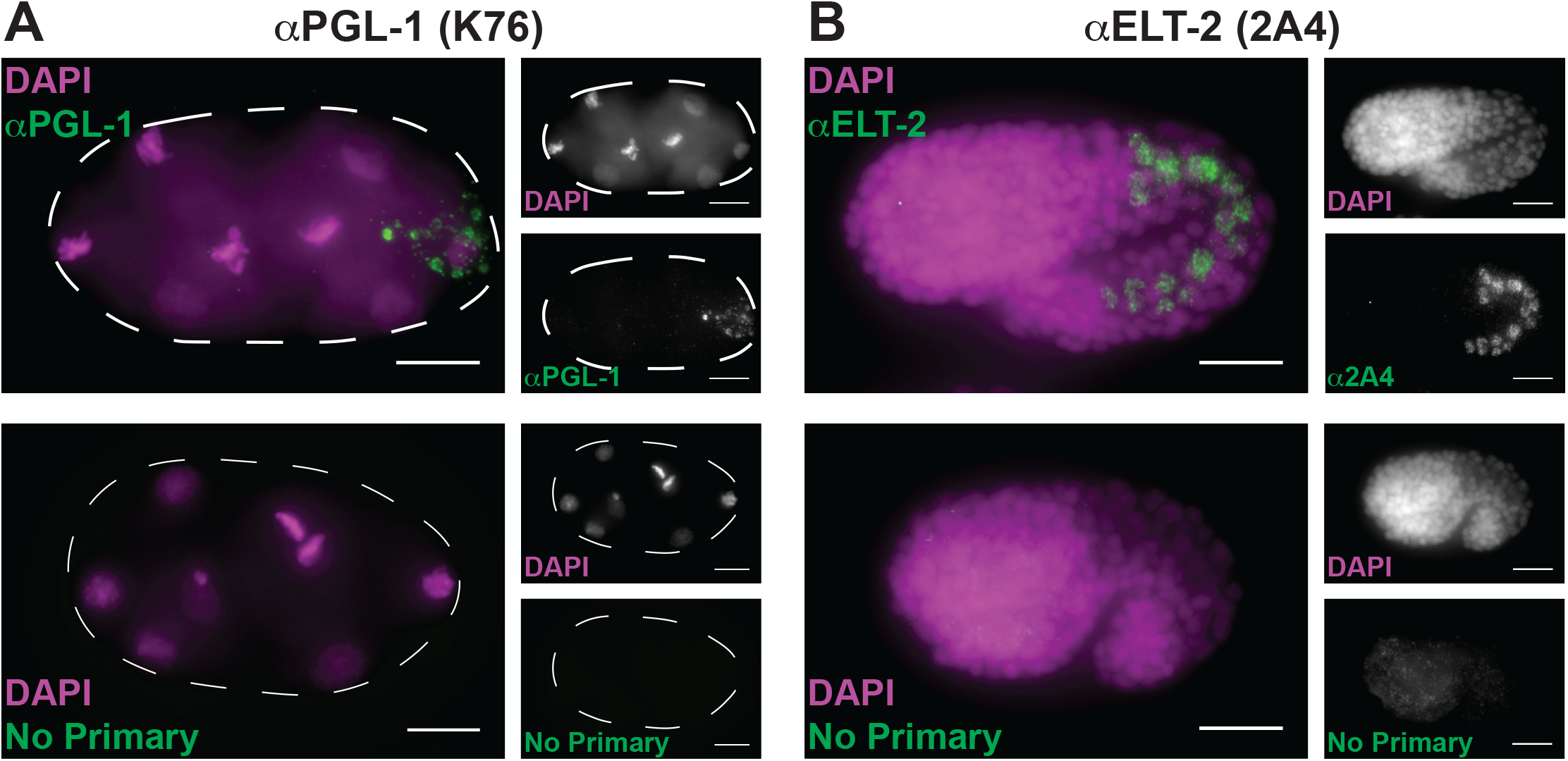
Simplified immunofluorescence in *C. elegans* embryos. Immunofluorescence was performed on N2 embryos as described (Protocol 3). Embryos were incubated with 1:20 dilutions of K76 (DHSB, Antibody registry ID AB_531836) (**A)** or 1:1000 dilutions 2A4 (DHSB, Antibody Registry ID: AB_2618114) **(B)** primary antibodies followed by incubation with 1:250 dilutions ofAlexa Fluor Goat Anti-Mouse secondary antibody (Jackson ImmunoResearch, Antibody Registry ID: AB_2338840) (green). In the presence of K76 (anti-PGL-1), P granules are observed **(A, top)**, while 2A4 (anti-ELT-2) stained the intestine-specific ELT-2 transcription factor **(B, top)**. Non-specific binding of the secondary was not observed in either instance **(A, B, bottom)**. Three biological replicates were performed for each experiment. Scale bars represent 10 μm.

### 2.3 smFISH and smiFISH

Single-molecule RNA Fluorsecence in situ Hybridization (smFISH) has provided insights into the regulation of transcripts in *C. elegans* at all stages of development. smFISH probes can be designed and synthesized in the lab^8,9^ or ordered as a set from Biosearch Technologies (Novato, CA). Some typical fluorophores include Cy5, Quasar 670, Alexa 594, Cal Fluor 610, and Fluorescein, among many others. In general, we have had the best signal to noise and most photostable fluorescence using Quasar 670 and Cal Fluor 610, which also work well in experiments probing for two RNAs. Fluorescein tends to have very low signal-to-noise ratios.

Because each probe in a set requires chemically conjugation with fluorophores for each specific transcript to be imaged, smFISH probe sets are relatively expensive^6,7,10^. Targeting a single RNA typically costs in the range of ∼$500. Recently, Tsanov et al. outlined a straightforward, flexible method for reducing the cost of single-molecule RNA detection: single-molecule inexpensive Fluorescence In Situ Hybridization (smiFISH). smiFISH brings down the cost of single molecule RNA detection by taking advantage of a single, universal fluorophore-labeled secondary probe annealed *in vitro* to gene-specific primary probes (Figure 3A). Primary smiFISH probes contain two main parts facilitating efficacy and cost reduction: the gene-specific region complementary to the transcript of interest and the FLAP region complementary to the fluorescently-labeled secondary probe. *In situ*, the complementary region of the primary probes bind to the target RNA while it’s FLAP region is annealed to a fluorophore-labeled secondary FLAP probe. This regime significantly reduces the cost of single-molecule RNA visualization by eliminating the need to create chemically conjugated probe sets for each specific target RNA. To test whether smiFISH performs as well as traditional smFISH in *C. elegans* embryos, we compared *nos-2* or *imb-*2 smFISH and smiFISH probes in the same sample (FIGURE 3). We found that smiFISH faithfully reproduces the sensitivity, spatial resolution, and reliability of smFISH probes. We have found that in larval stages smiFISH is less effective than smFISH using our standard protocols, possibly due to lower larval permeability preventing smiFISH probe entry.

**Figure 3.**
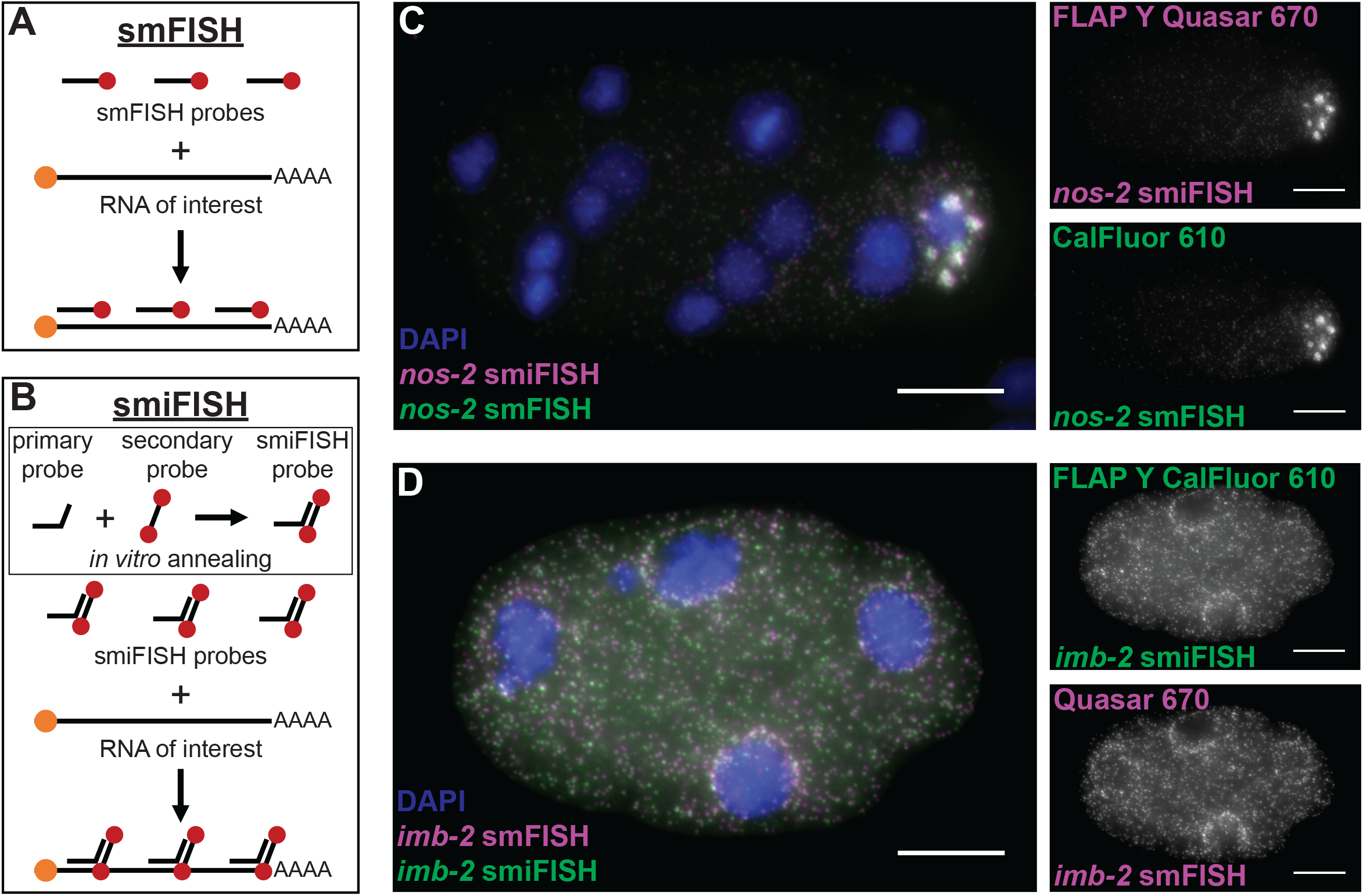
smFISH and smiFISH in *C. elegans* embryos. **A**. Schematic illustration of smFISH probes. **B**. Schematic illustration of smiFISH probes. **C**. *nos-2* RNA was visualized using smiFISH (magenta) and smFISH (green). *nos-2* smiFISH primary probes used FLAP-Y sequences and the secondary FLAP-Y probe was 5’ and 3’ dual-conjugated with Quasar 670 fluorophores. *nos-2* smFISH probes were 3’ single-conjugated with Cal Fluor 610. **D**. *imb-2* RNA was visualized using smFISH (magenta) and smiFISH (green). *imb-2* smFISH probes were 3’ single-conjugated with Quasar 670 fluorophores. *imb-2* smiFISH primary probes used FLAP-Y sequences and the secondary FLAP-Y probe was 5’ and 3’ dual-conjugated with Cal Fluor 610. Embryos were counterstained with DAPI in blue **(C, D)**. Three biological replicates were performed for each experiment using newly annealed smiFISH probes for each replicate. Scale bars represent 10 um.

### 2.4 smiFISH probe design

smiFISH primary probes can be designed as described Tsanov et al. 2016 using the R script Oligostan. Primary probes can be ordered in 96-well plates from IDT on the 25 nmol scale prediluted to 100 uM in IDTE buffer pH 8.0. Alternatively, if ordering 96 or more individual probes, oligos can be ordered on the 500 pm scale, which still provides ample primary probes for hundreds of experiments. For most experiments, ∼12-16 primary probes per transcript is sufficient, although testing as few as 8 primary probes has produced discernable single-molecule spots in *C. elegans* embryos. An increased number of primary probes typically increases the signal-to-noise ratio for any given transcript. Secondary FLAP probes (see smiFISH below) can also be ordered as 5’ and/or 3’ single- or dual-fluorophore-labeled oligos from either Biosearch Technologies or IDT (Coralville, Iowa).

### 2.5 Optimizing signal-to-noise in smFISH and smiFISH samples

In RNA FISH experiments, it is crucial to obtain the highest possible signal-to-noise ratio (SNR) to ensure reliable interpretation of the data. One common question surrounding smFISH is whether commercial reagents (i.e., Stellaris) are superior to homemade reagents^7,9^. By comparing the signal-to-noise ratio of four transcripts imaged by smFISH using homemade buffers or Stellaris buffers, we found Stellaris buffers perform significantly better for all four transcripts, ranging from 15-25% improvement in average SNR compared with homemade buffers. **(FIGURE 4)**. Another common concern with smFISH experiments is photolability. Due to the relatively low signal, high laser powers, and small number of fluorophores (∼24-48) utilized in smFISH experiments, photobleaching can occur rapidly. Photobleaching is of particular concern with thick samples that must be imaged through many Z stacks, as is the case with *C. elegans* embryos (∼12-20 um thickness as prepared in Protocol 1: 3.1.4, or ∼60-100 stacks per embryo at 0.2 um spacing between z-stacks). One of the primary causes of photobleaching is degradation of fluorophore molecules by oxygen radicals produced upon laser excitation^23^. Therefore, free-radical scavenging antifades are commonly used to reduce the degree of experimentally-induced photobleaching. We tested combinations of antifades to determine the optimal reagents for maintaining high signal-to-noise throughout an experiment. Through these experiments, we found that the optimal antifade solution can vary depending on the probe set or fluorophore **(FIGURE 5)**. In our hands, vectashield, N-propyl gallate, or a mixture of the two, provided the best signal stability for Cal Fluor 610 and Quasar 670 labeled RNAs in *C. elegans* embryos.

**Figure 4.**
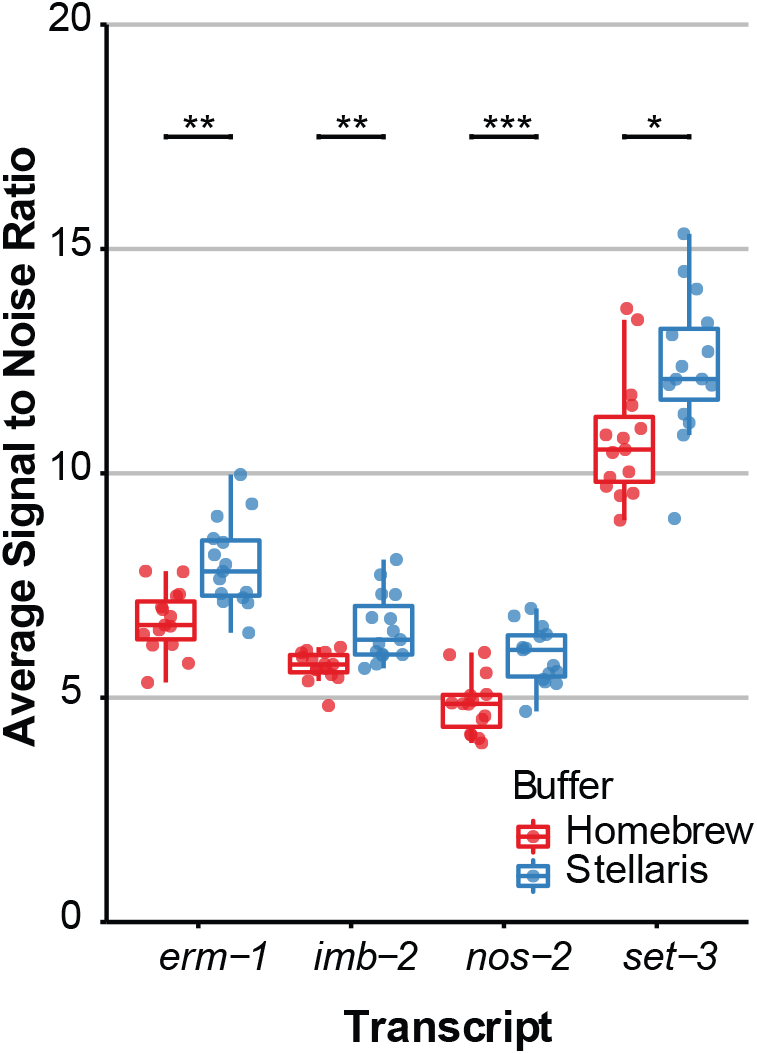
Stellaris buffers provide higher signal-to-noise ratios than homebrew buffers. Signal-to-noise ratios were calculated for each RNA puncta identified when smFISH was performed using homebrew (red) or commercial Stellaris (blue) buffers. The signal-to-noise ratio was calculated by identifying RNA spots using FISHquant^28^ before using the ImJoy SNR plugin (REF if published). In short, the SNR plugin compares the intensity at the coordinates of RNA puncta identified by FISHquant to the average intensity of a sphere surrounding the spot to calculate SNR. Four Stellaris smFISH probe sets were used, *erm-1* conjugated to Cal Fluor 610, *imb-2* conjugated to Quasar 670, *nos-2* conjugated to Quasar 670, and *set-3* conjugated to Cal Fluor 610. Individual dots represent the average SNR in one embryo. Three biological replicates were performed for each experiment, and 15 embryos were quantified for each condition. P values from Benjamini-Hochberg corrected t-tests are shown (0.05 > * > 0.005 > ** > 0.0005 > *** > 0.00005).

**Figure 5.**
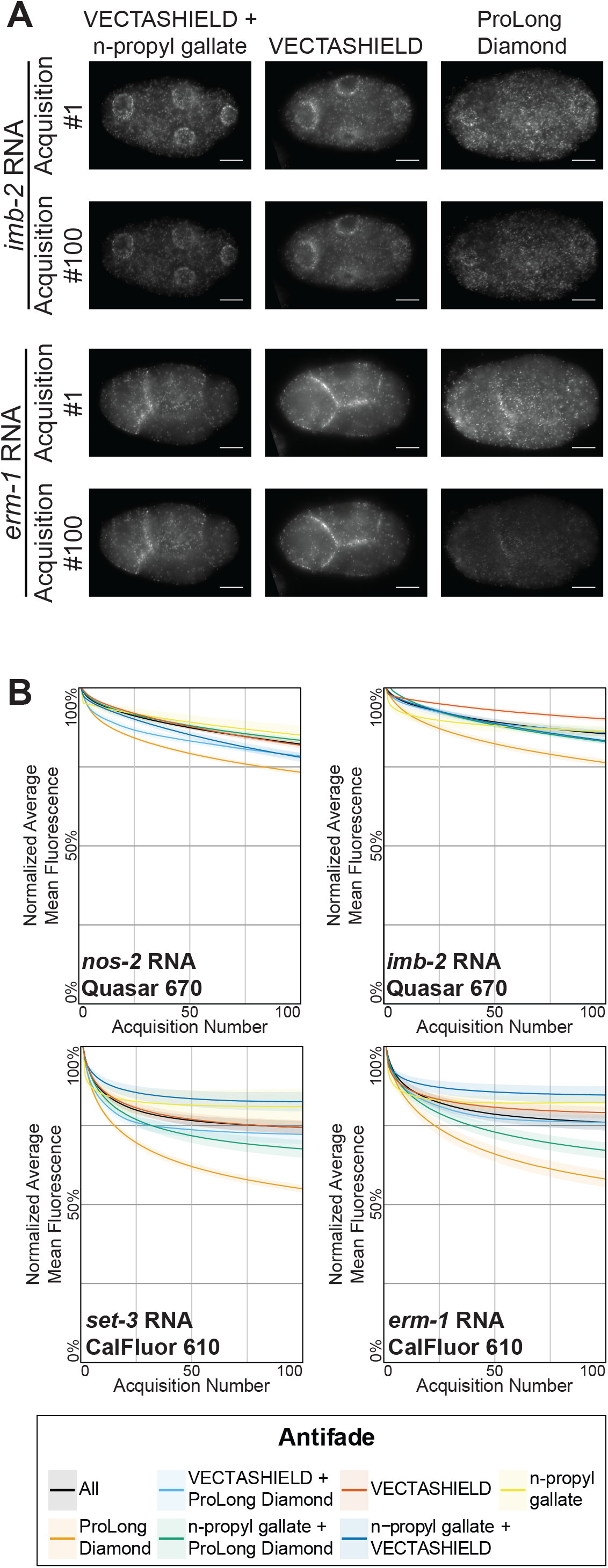
Effect of anti-fade composition on smFISH signal intensity. The mean fluorescence intensity of smFISH signal over 100 exposures was measured in embryos using various antifades and their combinations. Experiments were performed using four different smFISH probe sets: *erm-1* conjugated to Cal Fluor 610, *imb-2* conjugated to Quasar 670, *nos-2* conjugated to Quasar 670, and *set-3* conjugated to Cal Fluor 610). **A**. The average mean intensity throughout imaging was normalized to the intensity of first acquisition for each embryo. The shaded region represents the standard error of the mean for each exposure. Three biological replicates were performed for each experiment, and no less than nine embryos were quantified for each condition. **B**. Representative images of the first and final acquisitions for *imb-2* (top) and *erm-1* (bottom) RNAs using VECTASHIELD and N-propyl gallate (left), VECTASHIELD only (middle), and ProLong Diamond (right) anti-fades.

### 2.6 Sequential IF/FISH protocol

Simultaneous detection of an RNA and its cognate protein reveals a wealth of information regarding the expression patterns, regulation, and functions of genes. However, the combination of IF and FISH is often challenging due to slight incompatibilities in traditional protocols. Typically combined IF/FISH protocols require specific tailoring to the system of interest^24–26^. This includes one protocol designed for the extruded *C. elegans* gonad, which requires hand dissection of individual animals and careful slide preparation^27^. When immunofluorescence is performed in series with smFISH all reagents must be RNAse free where possible. Steps containing BSA must be treated with an RNAse inhibitor to prevent RNA degradation. We demonstrate a sequential IF/FISH protocol using the anti-PGL-1 antibody, K76 and smFISH probes against the P granule RNAs *nos-2* (Figure 6A) and *cpg-2* (Figure 6B). Additionally we show IF/FISH results in embryos stained with the ELT-2 antibody, 2A4 and hybridized with smFISH probes targeting *elt-2* RNA (Figure 6C)

**Figure 6.**
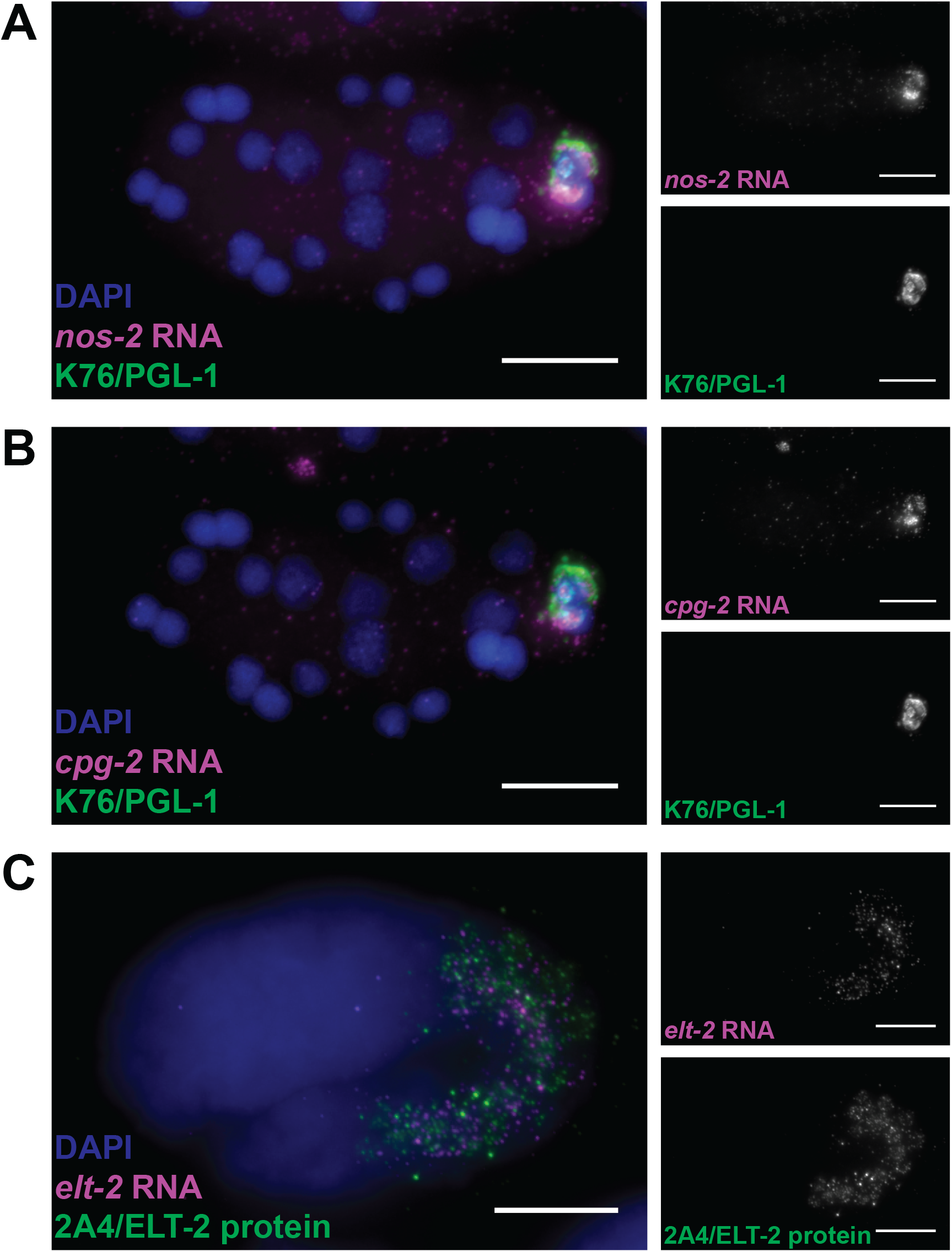
Sequential IF/FISH. Immunofluorescence followed by smFISH was performed on N2 embryos. IF was performed using K76 (**A and B)** or 2A4 **(C)** primary antibodies to identify PGL-1 containing P granules and ELT-2 protein (magenta), respectively. smFISH was used to simultaneously detect the P granule constituent RNAs *nos-2* (**A)** and *cpg-2* **(B)**, or *elt-2* mRNA **(C)**, all in magenta. Embryos were counterstained with DAPI (blue). Three biological replicates were performed for each experiment. Scale bars represent 10 μm.

### 2.7 Simultaneous IF/FISH protcol

If performing IF with a high-affinity nanobody or single chain variable fragment (ScFv), a simplified protocol can often be utilized. Under these circumstances, the FISH protocol (Protocol 3) can be followed with the caveat that fluorescently labeled nanobody or ScFv can be added directly to the hybridization buffer in step 4 and incubated with the FISH probes and sample overnight to perform IF. It is unclear why some nanobodies and ScFv work with this simplified protocol, but it is possible that their small size compared to traditional antibodies allows better permeation during hybridization while the high-affinity of some common nanobodies/ScFv facilitate antigen recognition at the higher temperatures required for RNA FISH probe hybridization. Here we present results for simulataneous IF/FISH from embryos containing PATR-1::GFP (Figure 7). The embryos were stained with a Janelia Fluor 549 (Tocris cat. no. 6147) labeled anti-GFP nanobody (Chromotek, gt-250) in hybridization buffer along with smFISH probes targeting *nos-2* RNA.

**Figure 7.**
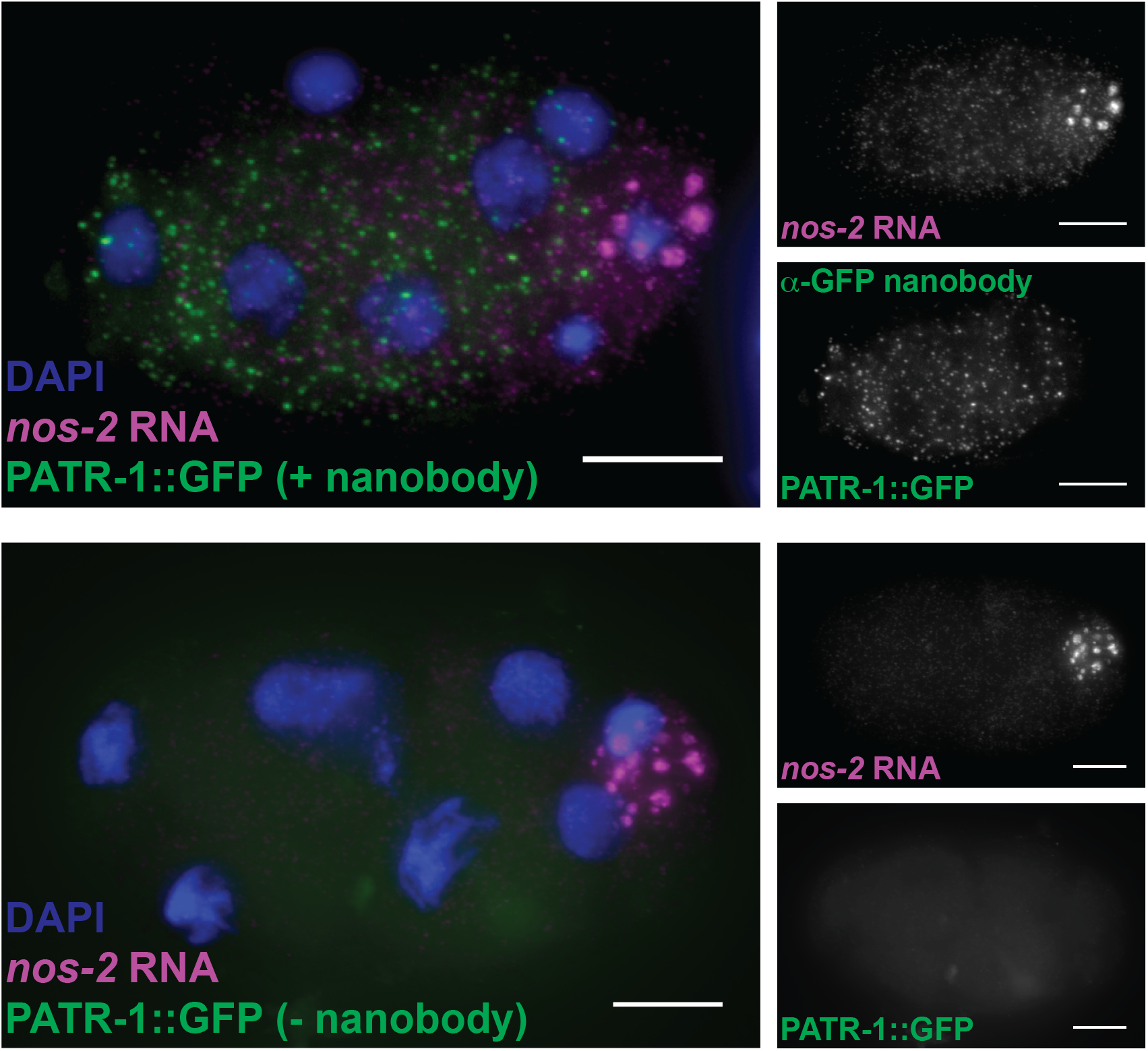
Simultaneous IF/FISH. smFISH was performed on N2 embryos with the addition of anti-GFP nanobody to hybridization buffer. *nos-2* mRNA (magenta) was probed using smFISH probes conjugated to Quasar 670. PATR-1::GFP (green) signal was visualized using 2.37 ug/ml Janelia Fluor 549 (Tocris 6147) conjugated anti-GFP nanobody (Chromotek, gt-250) (top). A no nanobody control is also shown (bottom). DNA was counterstained with DAPI (blue). Three biological replicates were performed for each experiment. Scale bars represent 10 μm.

### 2.8 smFISH and smiFISH data analysis

Depending on the biological questions at hand, there are several routes for the interpretation of smFISH data. These analyses range from simply characterizing the quality of the data, counting the number of RNAs in the samples, or even identifying spatial distributions of RNA within cells of interest.

The most common method for quantification of smFISH data is counting the number of RNAs within the sample. Some commonly used tools for this purpose are FISH-quant^28^ and StarSearch^8^. These algorithms function by enhancing spot signals through various filtering methods, setting a threshold for RNA spot detection, and identifying individual spots. Thresholds are often set manually by testing a range of intensity values. When plotting these values against the number of detected spots, a plateau can often be seen corresponding to threshold values separating RNA spots from lower intensity noise. When performing spot detection analysis of smFISH data, it is imperative to ensure the SNR of the data is sufficient to identify spots unambiguously. SNR can be calculated using an ImJoy plugin, which compares the intensity of a detected spot to the surrounding background intensities (https://github.com/fish-quant). In our experience, if SNR values are below ∼3-4, spot detection becomes less reliable. When analyzing smFISH data using FISH-quant or StarSearch, if there is no clear plateau of RNA counts over various threshold values, the SNR is likely too low for accurate RNA spot detection.

As smFISH has become more widely utilized, novel methods of analysis beyond spot counting are rapidly developing. For instance, FISH-quant has been ported from Matlab to an open-source implementation in Python and successfully applied to two large-scale screen projects^29,30^. This package includes methods for detecting, deconvolving overlapping RNAs to increase the counting accuracy of highly abundant or clustered RNAs^5,30^, measuring the signal-to-noise ratio of an image (https://github.com/fish-quant), and even identifying diverse subcellular localization patterns of RNA^30,31^. Further, to facilitate its usage by non-specialists, several plugins providing user-interfaces for the data analysis platform ImJoy^32^ were developed. As more labs adopt smFISH methodologies and more high-throughput methods of in situ RNA detection develop^33– 37^, more sophisticated analysis methods are likely to arise. An exciting initiative is Starfish, an open-source software suite with the goal to build a unified data-analysis tool and file format for several spatial transcriptomic techniques^38^.

### 2.9 IF data analysis

Standard methods of analysis for IF experiments include measuring the total internal fluorescence and measuring colocalization between different markers. These methods require that imaging conditions, such as laser intensity and exposure times, are held constant across samples and replicates. We will highlight publicly available tools for analysis here; however, most microscopes ship with instrument-specific software packages capable of performing these analyses. Total internal fluorescence compares the intensity of a protein visualized by IF in a control sample and an experimental condition, such as an RNAi knockdown or protein knockout. Total internal fluorescence can be measured over the total volume of the embryo, or regions of interest can be masked either automatically or manually if specific regions must be analyzed. Regardless of whether particular segmentations are required, these analyses can be performed relatively quickly in FIJI Is Just ImageJ (FIJI)^39,40^. Additionally, several FIJI plugins are available to analyze a protein of interest’s colocalization with another fluorescent marker. It is crucial when performing colocalization analyses to consider optimal uses for any given colocalization metric, as there are well-documented circumstances where these metrics can be misleading^41^. Helpful instructions for segmentation, colocalization analysis, and much more can be found at https://imagej.net/.

### 2.10 Combined IF/FISH data analysis

As with the analysis of IF data, colocalization analyses may be performed on combined IF/FISH data. However, due to the punctate nature of FISH signal, RNA spots may not overlap with a colocalization marker as well as expected, resulting in deceptively low colocalization coefficients. This can occur for several reasons. First, the small total volume of RNA puncta can lead to high variability in colocalization. This variability is compounded by the low temporal resolution of fixed cell experiments and the stochastic movements of RNA in the cell, even for tightly localized transcripts. Moreover, because it is often not known what proteins an RNA may be directly interacting with, it can be more desirable to compare RNA distributions to a nearby landmark rather than an overlapping component. For these reasons, several groups are developing novel metrics for comparing RNA and protein data and analyzing the spatial relationships between them. For instance, by spatially modeling the coordinates of each RNA puncta and comparing their distributions to other RNAs or organelles, it is possible to identify RNA patterning at various cellular features such as cortical membranes, nuclear membranes, condensates/puncta, cellular protrusions, centrosomes, and more^5,29–31^.

## 3 PROCEDURES

### 3.1 Protocol 1: Sequential IF/smFISH Protocol (Embryo prep + fixation, immunofluorescence, smFISH, slide preparation)

*This protocol describes methods for isolating C. elegans embryos and fixing them in a manner compatible with both immunofluorescence and RNA FISH. Steps for performing immunofluorescence subsequently followed by smFISH are then outlined. Finally, slide preparation is described. This approach can be used for simultaneous visualization of RNA transcripts and a protein of interest in the same sample provided the FISH probes and fluorescent antibody are selected in distinct channels*.

#### 3.1.1 – Embryo Prep and Fixation

##### Reagents

1. 100% reagent grade acetone (Fisher cat. no. A18-500)
2. 100% reagent grade methanol (Fisher cat. no. A412-500)
3. Bleaching solution for use when imaging embryos (per 50 mL, make fresh):
  a. 40 mL deionized, distilled water
  b. 7.2 mL 5 M NaOH (Fisher cat. no. S318-400)
  c. 4.5 ml 5% NaHOCl (Ricca cat. no. 7495.5-32)
4. M9 buffer
  a. 3 g KH_2_PO_4_ (Sigma cat. no. P0662-500G)
  b. 6 g Na_2_HPO_4_ (Sigma cat. no. RDD022-500G)
  c. 5 g NaCl (Fisher cat. no. S271-500) Deionized, distilled water (ddH_2_O) to 1 L final volume Sterilize by autoclaving.
  d. Add 1 ml 1 M MgSO_4_ (Millipore cat. no. MX0075-1) using sterile technique after solution cools to prevent precipitation.

##### Embryo Preparation and Fixation Protocol

1. Grow worms to gravidity on OP50 seeded NGM plates. Synchronize by bleaching if necessary. We typically harvest one or two gravid 10 cm NGM plates seeded with ∼2 ml OP50 for each slide to be made. Other bacterial stocks, such as inducible RNAi vector containing *E. coli*, can be used if desired.
2. Harvest gravid worms by washing them off of plates using M9 and collect in a 15 mL conical tube in ∼15 ml total volume. Aggressive pipetting will increase yield by releasing more worms from the plates. Be sure not to pierce the plate’s surface as agar carried into the sample will persist.
3. Spin conical at 2000 x g for 1 minute to pellet gravid worms. Alternatively, allow gravid worms to settle over time.
4. Remove supernatant using a pipette or aspirator, being careful not to disturb worm pellet.
5. Resuspend worm pellet in 15 mL M9.
6. Spin to pellet again as above (3).
7. Repeat steps 4 - 6 until the supernatant is clear, removing supernatant after the final wash.
8. Add ∼15 mL of bleaching solution to the worms and nutate or hand-shake for 6-8 minutes until embryos are released from the mothers. Check on the condition of worms periodically throughout bleaching. The gravid adults should be broken into about two pieces before continuing. If worms are bleached for too long, some early-stage embryos may be damaged. For tips on harvesting embryos, see Porta-de-la-Riva et al. 2012^42^.
9. Centrifuge conicial at 2000 x g for 1 minute to pellet. Immediately remove supernatant and quench bleaching with 15 mL M9. At this point, embryos typically stick to the tube, and the supernatant can be carefully decanted to decrease the time before quenching.
10. After adding M9, vortex the pellet to release remaining worm fragments before centrifuging at 2000 x g for 1 minute.
11. Wash with 15 mL M9 two more times (for a total of 3 washes), vortexing the pellet after the addition of M9 each time. The aroma of bleach should be completely gone by the end of washing.
12. Transfer remaining embryos to a 1.7 ml microcentrifuge tube and pellet in a tabletop centrifuge for 30 seconds at 2000 x g. Turn tube 180° and repeat until a pellet has formed. Remove any remaining M9.
13. Add 1 mL pure methanol cooled to -20°C, vortex to break up the pellet, and immediately submerge in liquid nitrogen for 1 minute to crack the eggshell and promote permeabilization.
14. Remove the tube from liquid nitrogen and immediately begin pelleting at 2000 x g in 30 sec intervals, rotating the tube 180° between each spin. The sample will still be partially frozen for the first spins, but it is best to get the sample pelleting early to prevent over-fixation.
15. Once the embryos are pelleted, and the sample has been in methanol for 5 min, remove the methanol and replace it with 1 mL pure acetone cooled to -20°C. Store the sample at - 20°C for ∼3 min.
16. Pellet embryos by centrifugation as in step 14.
17. After embryos have fixed in acetone for 5 min, remove the acetone and immediately continue to IF, smFISH, smiFISH, or IF/FISH protocol.

#### 3.1.2. Immunofluorescence

##### Reagents

1. 10X PBST
  a. 80 g NaCl (Fisher cat. no. S271-500)
  b. 2 g KCl (Sigma cat. no. P3911-500G)
  c. 14.2 g Na_2_HPO_4_ (Sigma cat. no. RDD022-500G)
  d. 2.4 g KH_2_PO_4_ (Sigma cat. no. P0662-500G)
  e. 1% Tween^®^ 20 detergent (w/v) (Sigma cat. no. P1379-500ML) Deionized, distilled water to 1 L final volume Sterilize by autoclaving Dilute to 1X in sterile deionized, distilled water
2. Bovine Serum Albumin (Sigma cat. no. A9418-5G)
  a. RNAse free BSA can be used if issues with RNA degradation occur with sequential IF/smFISH protocols; however, it is much more expensive.
3. Primary antibody or fluorescently labeled nanobody/ScFv
4. Fluorescent secondary antibody (if using an unlabeled primary antibody)
5. DAPI, 4’,6-Diamidino-2-Phenylindole, Dihydrochloride (Invitrogen cat. no. D1306)
6. RNasin^®^ Ribonuclease Inhibitor (If performing IF/FISH) (Promega cat. no. N2111)
7. 20X SSC (If performing IF/FISH)
  a. 800 ml deionized, distilled water
  b. 175.2 g NaCl (Fisher cat. no. S271-500)
  c. 88.2 g sodium citrate tribasic dihydrate (Sigma cat. no. S4641-500G) pH to 7.0 with 1 M HCl. Deionized, distilled water to 1 L and autoclave. Dilute to 2X in sterile deionized, distilled water.

##### PRELIMINARY NOTES

If performing IF/FISH, all reagents must be RNAse free where possible. Steps containing BSA must be treated with an RNAse inhibitor to prevent RNA degradation (see step 6 and 8).

Once a fluorescent antibody has been added (either primary or secondary) all subsequent steps should be carried out in the dark, ie covered in foil, to minimize fluorophore bleaching.

##### Immunofluorescence Protocol

1. Prepare fixed embryo samples as described in 3.1.1 steps 1-17.
2. Add 1 ml 1X PBST to sample and nutate for 5 min to wash.
3. Pellet embryos by centrifuging at 2000 x g in 30 sec intervals, rotating the tube 180° between each spin until pellet forms.
4. Pipet or aspirate as much of the supernatant PBST as possible without disrupting the pellet.
5. Repeat steps 2-5 two more times (3 washes total).
6. Block for 30 min. at 37°C in 50-250 ul 1X PBST containing 1% w/v BSA with nutation. **IMPORTANT:** If FISH will be performed subsequently, it is essential to add 1 unit/ul RNasin^®^ (Promega) to prevent RNA degradation during steps where BSA is included.
7. Centrifuge embryos at 2000 x g in 30 sec intervals, rotating the tube 180° between each spin until pellet forms.
8. Pipet or aspirate as much of the supernatant as possible without disrupting the pellet.
9. Apply 25-100 ul 1° antibody diluted in 1X PBST with 1% w/v BSA (and 1 unit/ul RNasin^®^ if FISH will be performed subsequently). Nutate at room temperature for at least 1-2 hrs, or overnight at 4°C. Overnight incubations will give better IF signal, but can increase RNA degradation. Optimal antibody concentrations must be determined for each antibody.
10. Add 1 ml 1X PBST directly to sample and nutate for 5 min to wash out free antibody.
11. Centrifuge embryos at 2000 x g in 30 sec intervals, rotating the tube 180° between each spin until pellet forms.
12. Pipet or aspirate as much of the supernatant PBST as possible without disrupting the pellet.
13. Repeat steps 9-11 two more times (3 washes total).
14. Apply 25-250 ul fluorescently labeled 2° antibody diluted in 1X PBST and incubate for 1-2 hrs in the dark at room temperature with nutation. Optimal antibody concentrations must be determined for each antibody.
15. Add 1 ml 1X PBST and nutate for 5 min to wash out excess antibody.
16. Centrifuge embryos at 2000 x g in 30 sec intervals, rotating the tube 180° between each spin until pellet forms.
17. Pipet or aspirate as much of the supernatant PBST as possible without disrupting the pellet.
18. Repeat steps 15-17.
19. Add 1 ml 2X SSC and nutate for 5 min to equilibrate embyros in an smFISH compatible solution.
20. Centrifuge embryos at 2000 x g in 30 sec intervals, rotating the tube 180° between each spin until pellet forms.
21. Pipet or aspirate as much of the supernatant SSC as possible without disrupting the pellet.
22. Repeat steps 19-21.
23. Continue to 3.1.3, smFISH protocol

#### 3.1.3. smFISH

##### Reagents

1. Wash Buffer A (10% volume/volume formamide)
  a. 600 uL Stellaris Wash Buffer A (Biosearch Technologies cat. no. SMF-WA1-60)
  b. 2.1 mL DEPC treated RNAse free water (Invitrogen cat. no. AM9922)
  c. 300 uL deionized formamide (Millipore cat. no. S4117) Prepare 3 mL for each sample to be hybridized. Prepare Wash Buffer A fresh for each experiment.
2. Wash Buffer B
  a. Stellaris Wash Buffer B (Biosearch Technologies cat. no. SMF-WB1-20 Be sure to add 88 mL RNAse free water (Invitrogen cat. no. AM9922) to Wash Buffer B stock before use.
3. Hybridization Buffer (10% volume/volume formamide) Prepare 110 ul for each sample in an experiment Prepare hybridization buffer fresh for each experiment
  a. 99 uL Stellaris Hybridization Buffer (Biosearch Technologies cat. no. SMF-HB1-10/0
  b. 11 uL deionized formamide (Millipore cat. no. S4117)
4. Mounting Medium (5 mL)
  a. 2.5 mL 100% glycerol (Sigma cat. no. G5516-100ML)
  b. 100 mg N-propyl gallate (Sigma cat. no. 02370-100G)
  c. 400 ul 1 M Tris pH 8.0 (Sigma cat. no. 10708976001) N-propyl gallate is toxic. Vortex until N-propyl gallate has dissolved. Store mounting medium in amber tubes or covered in foil at either 4 or -20 °C. The solution is light sensitive. Throw mounting medium away if it begins to yellow or crystalize.
5. smFISH probes and/or annealed smiFISH probes
6. DAPI, 4’,6-Diamidino-2-Phenylindole, Dihydrochloride (Invitrogen cat. no. D1306)
7. RNAse free water (Invitrogen cat. no. AM9922)

##### smFISH Protocol

1. Prepare fresh buffers by adding formamide to Wash Buffer A and Hybridization Buffer. Wash Buffer A and Hybridization Buffer should always have formamide added immediately preceding the experiment. Formamide can decompose over time, particularly at higher temperatures, leading to less stringent probe binding. It can also acidify when exposed to air resulting in fluorophore quenching. Formamide stocks should be stored frozen and their pH monitored periodically (pH 7-8 is ideal)
2. Add 2 ul 1.25 uM smFISH probes (1:20 dilution of 25 uM stocks) to 110 ul hybridization buffer. If performing experiments using multiple probe sets with different fluorophores, add 2 uL of each probe set. Mix well. Hybridization buffer is viscous. Optional step: If performing Protocol 4 (simultaneous IF/FISH) using a compatible ScFv or nanobody, additionally add the appropriate concentration of ScFv or nanobody to the hybridization buffer. Note: Although 2 uL has worked well for most of the probe sets we have used, it is helpful to perform a titration over ∼1 order of magnitude of concentrations to identify optimal probe concentrations on an individual probe set basis.
3. Centrifuge embryos at 2000 x g in 30 sec intervals, rotating the tube 180° between each spin until pellet forms.
4. Pipet or aspirate as much supernatant as possible without disturbing the pellet.
5. Prehybridize sample in 1 mL Wash Buffer A and incubate at room temperature for ∼5 minutes.
6. Centrifuge embryos at 2000 x g in 30 sec intervals, rotating the tube 180° between each spin until pellet forms.
7. Pipet or aspirate as much supernatant as possible without disturbing the pellet.
8. Add 100 uL hybridization buffer with probes to the pelleted embryos and hybridize at 37 °C in the dark for 8-48 hours. Store prepared Wash Buffer A at room temperature or 37°C during this incubation. Warm buffer will increase the stringency of probe binding and decrease background and non-specific binding. If available, use a thermomixer to shake the hybridization solution and all subsequent washes at 450 rpm during incubation to ensure even probe penetration.
9. Add 1 mL Wash Buffer A directly to the embryos in hybridization solution.
10. Incubate at 37°C in the dark for 30 minutes.
11. Centrifuge embryos at 2000 x g in 30 sec intervals, rotating the tube 180° between each spin until pellet forms.
12. Pipet or aspirate as much supernatant as possible without disturbing the pellet.
13. Add 1 mL Wash Buffer A containing 1 ng/uL DAPI to the sample.
14. Incubate at 37°C in the dark for 30 minutes.
15. Centrifuge embryos at 2000 x g in 30 sec intervals, rotating the tube 180° between each spin until pellet forms.
16. Pipet or aspirate as much supernatant as possible without disturbing the pellet.
17. Add 1 mL Wash Buffer B and incubate for ∼5 minutes.
18. Repeat step 15 and 16.
19. Resuspend in 50 uL of mounting medium (or less if the sample is small) and incubate at 4 °C for 30 minutes to ensure antifade penetrance.
20. Move to slide preparation.

#### 3.1.4 Slide Preparation

##### Reagents

1. VECTASHIELD mounting medium (Vector Laboratories cat. no. H-1000-10)
2. 8mm 1.5 thickness round cover glass (Electron Microscopy Sciences, cat. no. 72296-08)
3. Glass microscope slides (VWR cat. no. 48312-401)
4. 1.5 thickness, 22×22 mm coverglass (VWR cat. no. 48366-227) Use the appropriate thickness for your microscope.
5. Grace Bio-Lab Press-To-Seal silicon isolator (Sigma cat. no. GBL664504-25ea)

##### Slide Preparation Protocol

1. Working at a dissecting microscope, drop 2 – 6 ul of embryos suspended in mounting medium onto a single 8 mm 1.5 thickness round cover glass resting on a glass slide. Always wear gloves when handling slides and cover slips to prevent smudging and contamination.
2. Add the same volume of VectaShield VECTASHIELD antifade solution and pipet up and down to mix thoroughly. Try to keep the final volume to ∼4-6 ul by removing some of the mixture. This is a good time to break up any large clumps of embryos by pipetting.
3. Place a 1.5 thickness 22 mm x 22 mm square cover glass on top trying to avoid bubbles. Do not let the coverslip touch the slide. The sample solution will pour over the edge of the round coverslip and seal it to the slide beneath through surface tension. Having the round coverslip close to the edge of the slide can provide some extra working height. Additionally, gently lowering the square coverslip from front to back over the round coverslip until surface tension pulls the round cover slip up will help prevent spillover.
4. Flip the coverslips so the square coverslip is on the bottom. Remove as much liquid as possible from between the two cover glasses using a torn kimwipe placed against the round one. The aim is to flatten the embryos as much as possible without damaging them. Samples can be firmly pressed on with a pipette tip as long as the coverslip doesn’t slide from side to side. The ideal depth of an embryo on the slide is ∼12-20 um. Signal-to-noise ratio will decrease and photobleaching will increase with increasing thickness due to out-of-focus light and more image acquisitions, respectively.
5. Affix the cover slip sandwich to a microscope slide using a Grace Bio-Lab Press-To-Seal silicon isolator such that the embryos will be imaged through the square coverslip.
6. Head off to the microscope!

### 3.2 Protocol 2: smFISH or smiFISH alone (Embryo prep + fixation, smFISH or smiFISH, slide preparation)

*This protocol describes the workflow for performing smFISH or smiFISH in embryos, from sample preparation to slide preparation*.

#### 3.2.1 Embryo Prep and Fixation

- Perform Embryo prep and fixation as in 3.1.1

#### 3.2.2 smFISH

- Perform smFISH as in 3.1.3

#### 3.2.3. smiFISH

1. Perform smiFISH as in 3.1.3 with the following considerations/exceptions
  - The following reagents and protocol is required to generate annealed primary + secondary smiFISH probes. <u>Reagents</u>
    1. 8-24 gene specific primary probes resuspended at 100 uM in IDTE pH 8.0 (or Tris pH 8.0)
    2. 1 Fluorophore-labeled FLAP probe resuspended at 50 uM in Tris pH. 8.0
    3. New England Bio Labs Buffer 3 (or 3.1) (NEB cat. no. B7203S) <u>smiFISH probe annealing</u>:
      i. Combine primary probes at equimolar ratio and dilute to 0.833 uM in Tris pH 8.0. This primary probe mixture is stable at -20°C indefinitely. In a PCR tube, prepare a solution of:
      ii. 2 uL primary probe set
      iii. 1 uL 50 uM FLAP secondary probe
      iv. 1 uL NEB Buffer 3 (or 3.1)
      v. 6 uL RNAse free water Anneal primary probe set to fluorophore-labeled secondary probes using the following thermocycling conditions:
      vi. 1 cycle at 85 °C for 3 minutes
      vii. 1 cycle at 65 °C for 3 minutes
      viii. 1 cycle at 25 °C for 5 minutes Annealed smiFISH probes are viable at -20 °C for up to at least a week. Treat annealed smiFISH probes as diluted smFISH probes. 2 ul annealed smiFISH probe works well for most hybridizations

#### 3.2.4 Slide Preparation

- Prepare slides as in 3.1.4

### 3.3 Protocol 3: Immunofluorescence alone (Embryo prep + fixation, immunofluorescence, slide preparation)

*This protocol describes the steps to perform immunofluorescence in C. elegans embryos from harvesting embryos to preparing slides*.

#### 3.3.1 Embryo Prep and Fixation

- Perform Embryo prep and fixation as in 3.1.1

#### 3.3.2. Immunofluorescence

- Perform immunofluorescence as in 3.1.2 with the following exceptions:
  1. At step 15, nutate the sample in 1X PBST for 10 minutes instead of 5.
  2. Pellet embryos by centrifuging at 2000 x g in 30 sec intervals, rotating the tube 180° between each spin until pellet forms.
  3. Pipet or aspirate as much of the supernatant PBST as possible without disrupting the pellet.
  4. Counterstain with 1X PBST containing 2 ul 500 ng/mL DAPI for 10 min.
  5. Pellet embryos by centrifuging at 2000 x g in 30 sec intervals, rotating the tube 180° between each spin until pellet forms.
  6. Pipet or aspirate as much of the supernatant PBST as possible without disrupting the pellet.
  7. Add 1 ml 1X PBST directly to sample and nutate for 10 min to wash out excess DAPI.
  8. Repeat steps 5-7, followed by steps 5 and 6 (for two 1X PBST washes).
  9. Resuspend in 50 uL of mounting medium (or less if the sample is small) and incubate at 4 °C for 30 minutes to ensure antifade penetrance.

#### 3.3.3. Slide preparation

- Prepare slides as in 3.1.4

### 3.3 Protocol 4: Abreviated protocol for IF/smiFISH for use with nanobodies. (Embryo prep + fixation, simultaneous IF/smiFISH, slide preparation)

*This protocol describes a simplified method for performing immunofluorescence at the same time as smFISH with select antibodies*

#### 3.3.1 Embryo Prep and Fixation

- Perform Embryo prep and fixation as in 3.1.1.

#### 3.3.2. Simultaneous immunofluorescence and smFISH

- Perform smFISH as in 3.1.3 with the following exceptions and considerations:
  - This protocol only works with a subset of antibodies.
    ▪ We have had the best results using high-affinity nanobodies, ScFv, or fragmented antibodies^43^. High-affinity, small sized antibodies have improved the success of this simplified protocol in our hands.
    ▪ We have only had success with primary staining using this protocol. Immunofluorescence using secondary antibody amplification during wash steps has not succeded.
  - At step 2, when preparing the hybridization buffer mix, incorporate the appropriate concentration of antibody and proceed normally.

#### 3.3.3. Imaging & Analysis

- Perform imaging & analysis as in 3.1.4 with the following considerations/exceptions

## 4 CONTROLS AND TROUBLESHOOTING

### Validating new probe sets

There are several ways to validate new probe sets for target specificity and labeling efficiency. The most straightforward test for target specificity is to use the probes in a wildtype and deletion strain for the target of interest to ensure the probeset is binding only when the RNA is present. If a deletion allele is not available, RNAi can be utilized to a similar end. However, it is important to note that residual fluorescent signal may be present after RNAi because the knockdown may be incomplete or may only partially degrade the targets. Target specificity can also be determined by targeting a gene with two separate probe sets in different colors, which should colocalize if the probes are specific. Labeling efficiency of a probe set can be determined by comparing transcript abundance found using smFISH data to other sources, such as qRT-PCR, digital-droplet PCR, or quantitative sequencing data.

### Positive controls

Positive control smFISH probesets should be consistently employed to ensure the protocol is working. These probe sets have the added benefit of marking specific cell lineages or developmental stages and thereby identify the embryo’s orientation or stage. By comparing the performance across replicates, researchers can identify outliers or problems in protocol execution. When troubleshooting, the use of smFISH probe sets that anneal to highly abundant RNAs, such as the polyA sequence of mRNA, or using previously validated probes can be useful to ensure the FISH protocol is successful.

### Photobleaching

Due to the small number of fluorophores on any single RNA, the photolabile nature of common fluorophores, and the common use of widefield microscopy for FISH experiments, FISH can often suffer from rapid photobleaching. If a sample has clear puncta that disappear throughout imaging or the mean intensity of the sample drops rapidly during acquisition, photobleaching is likely reducing the data’s quality. Anti-fade should always be included in slide preparation and given time to permeate the sample before imaging to prevent photobleaching. Further, imaging from long, low energy wavelength lasers to short, higher-energy (i.e., from far-red to UV) can help preserve fluorescence.

### Low Signal to Noise

Since *C. elegans* embryos are relatively thick (∼20-30 um), the use of widefield microscopy will capture a large amount of out-of-focus signals from non-focal z-planes in the sample. Embryos can be flattened during slide preparation to improve SNR. We have found that samples from ∼12-20 um thick have an optimal signal-to-noise ratio without obviously perturbing sample morphology. While pressing down on embryos does not seem to affect their morphology, any lateral motion during slide preparation will shear embryos, so it is essential to press directly down when making slides.

### Crosstalk of smiFISH secondary probes

Tsanov et al. demonstrated that multiple primary probe sets containing the same FLAP sequence could be utilized in the same experiment without observable mislabeling by annealing them to secondary probes labeled with distinct fluorophores (i.e., probeset-1 FLAP-Y-Cal Fluor 610, probeset-2 FLAP-Y-Quasar 670). We have validated this in the *C. elegans* embryo.

### Probing for short transcripts

If a transcript is too short to design ample FISH probes, it can be worrisome to order probesets. We have obtained clear punctate signal for probe sets using as few as eight smiFISH probes. If a transcript is too short for even eight probes, it is worth considering amplification-based FISH methods^37,44–46^, which have been utilized in *C. elegans*^47^. However, quantification of amplification-based FISH is far less accurate due to variability in signal strength from single RNA molecules.

### smiFISH secondary aggregates

In some instances, we and other groups (personal comm) have observed large aggregates of fluorescently labeled secondary smiFISH probes on the surface of cells or adhered to slides. In our experience, vortexing annealed smiFISH probes followed by a quick centrifugation in a microfuge before hybridization and vigorous vortexing of samples after hybridization are sufficient to remove these large aggregates.

### Validation of antibodies

With any IF experiment, it is essential to validate the antibodies’ function and specificity. Primary antibodies can be validated using null or RNAi strains to ensure that the antibody is binding specifically to the target antigen. Secondary antibodies can be tested for specificity by incubating them in the absence of primary antibodies to ensure that there is no staining of endogenous antigens. Should an antibody have some non-specific binding, it may be possible to increase specificity by depleting the antibody using a null allele^11^. It is also necessary to test every antibody’s sensitivity over a range of concentrations to identify the optimal concentration for detecting the antigen of interest without promoting non-specific staining, typically over at least one to two orders of magnitude. Most commercial antibodies have a range of suggested optimal concentrations for immunofluorescence that can be used as a starting point. It is wise to test these concentrations for each experiment or experimental condition because changes in protein concentration or antigen accessibility can lead to different optimal concentrations of antibodies on a case-by-case basis. It is important to be aware that this can make downstream quantification inaccurate; however, so it is beneficial to use identical staining conditions when possible.

### Low yield

If embryo yield is low after performing IF, ensure that detergent is being used in the wash steps as it strongly reduces adherence to pipette tips and plastic tubes .

### Positive controls

If a protein can not be detected using a validated antibody, it is crucial to ensure that IF is working correctly. Staining common cytoskeletal components such as actin or microtubules can both verify the efficacy of the IF protocol in a sample while simultaneously demonstrating the sample is morphologically intact. Alternatively, a fluorescent protein, such as GFP, can be targeted for immunofluorescence using a different color and colocalization analyzed to ensure effective staining.

### RNA degradation

The most common issue in performing combined IF/FISH is RNA degradation. It is essential to use RNase-free reagents throughout the protocol and, when necessary, to add RNase inhibitors such as RNasin. In our experience, RNase inhibitor was only necessary during steps where BSA is present (which contains RNases). However, if RNA is not visible after performing IF/FISH, it is likely due to RNase contamination. Remaking reagents with RNase-free components or adding RNase inhibitors at each step will likely remedy this issue. As RNase inhibitor is relatively expensive, it is best to ensure the purity of reagents where possible. If RNA degradation continues to be an issue, reducing the duration of the IF steps of the protocol tends to improve RNA signal at the cost of protein signal. For example, performing a two-hour incubation with primary antibody instead of overnight can reduce RNA degradation.

### Permeabilization and fixation

*C. elegans* embryos are highly effective at preventing environmental contaminants from entering. This is in part due to the permeability barrier, a membranous barrier that prevents fluid exchange between the embryo and the environment^12^. The choice of fixative and fixation duration appear to be highly important for permeabilizing the embryo to antibodies, which are roughly 20X the mass and radius of smFISH probes (Ab ∼ 150 kDa and ∼ 60 A, 20mer oligo ∼ 7.5 kDa and ∼ 3 A^48,49^.). In our experience, a brief methanol fixation, liquid nitrogen freeze cracking, followed by a quick acetone fixation, was most effective at allowing antibodies to pass through the eggshell and permeability barrier while maintaining antigen recognition and FISH probe accessibility. The use of acetone was necessary for antibody staining. We interpret this result as acetone solubilizing permeability barrier components, thus increasing the size of molecules that can enter the embryo, although we have not rigorously examined the effective pore size under different fixation conditions. Our experiments with longer fixation times with both methanol and acetone reduced antigen recognition by antibodies (as well as GFP fluorescence for protein fusions). Moreover, the use of formalin/formaldehyde reduces the binding and photostability of FISH probes. Some antigens are likely more compatible with different fixatives, however. Should the fixation conditions presented here be incompatible with an antigen of interest, Duerr 2006^11^ describes alternative fixation strategies. If alternative fixation strategies must be pursued, it is crucial to keep in mind the effect they will have on the permeability of the eggshell and permeability barrier. If IF still fails, it may be worth using 150kDa fluorescent dextran to determine whether the embryo is permeable to antibodies.

### Clumps

For reasons unknown, in our experiments, *C. elegans* embryos that have undergone IF/FISH form aggregates of embryos that do not occur with either protocol alone. While some clumping seems inevitable, vigorous vortexing after fixation and every wash/pelleting step, as well as constant rocking during incubations, reduces the number and size of clumps. Clumps can also be disrupted by pipetting when preparing slides.

## ACKNOWLEDGEMENTS

We would like to thank Rob Williams and Wyatt Beyers for fruitful discussions about fixation methods and quantification strategies. We also thank Dustin Updike and James McGhee for reagents. A great thanks goes out to the entire ImJoy development team for making an invaluable platorm for data anlaysis. In particular, we extend our appreciation to Florian Mueller for help designing analysis pipelines, providing helpful insights, and feedback on the manuscript. Some strains were provided by the *Caenorhabditis* Genetics Center, which is funded by National Institutes of Health Office of Research Infrastructure Programs (P40 OD010440). Dylan Parker and Lindsay Winkenbach were supported in part by an NSF training grant (DGE-1450032).

## FUNDING

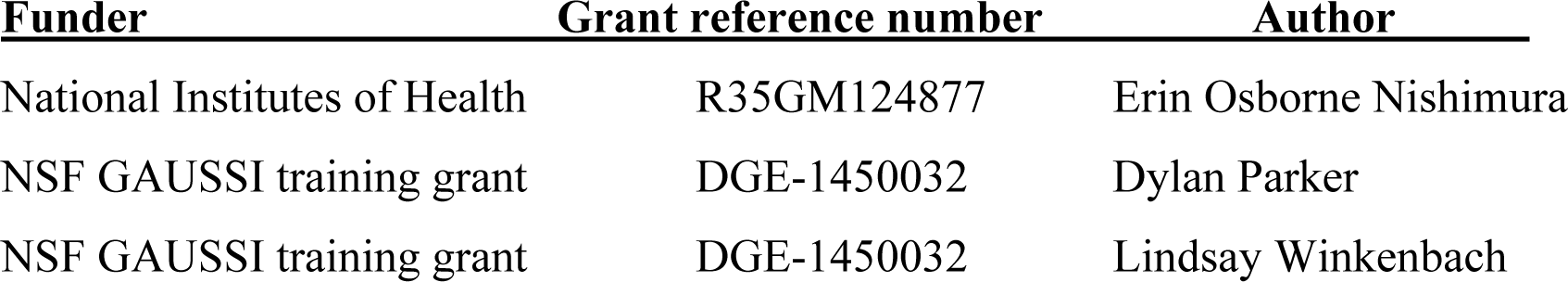

## AUTHOR CONTRIBUTIONS

Conceptualization: D.M.P., L.P.W., S.B., E.O.N.; Methodology: D.M.P., L.P.W., S.B., E.O.N.; Software: D.M.P., E.O.N.; Validation: D.M.P., L.P.W., S.B., E.O.N.; Formal analysis: D.M.P., A.P.; Investigation: D.M.P., L.P.W., S.B., A.P., E.O.N.; Resources: D.M.P., E.O.N.; Data curation: D.M.P., L.P.W., S.B., A.P.; Writing - original draft: D.M.P., E.O.N.; Writing – review & editing: D.M.P., L.P.W., S.B., A.P., E.O.N.; Visualization: D.M.P., L.P.W., S.B., E.O.N.; Supervision: D.M.P., E.O.N.; Project administration: D.M.P., E.O.N.; Funding acquisition: D.M.P., L.P.W., E.O.N.

